# Detecting past and ongoing natural selection among ethnically Tibetan women at high altitude in Nepal

**DOI:** 10.1101/223081

**Authors:** Choongwon Jeong, David B. Witonsky, Buddha Basnyat, Maniraj Neupane, Cynthia M. Beall, Geoff Childs, Sienna R. Craig, John Novembre, Anna Di Rienzo

## Abstract

Adaptive evolution in humans has rarely been characterized for its whole set of components, i.e. selective pressure, adaptive phenotype, beneficial alleles and realized fitness differential. We combined approaches for detecting selective sweeps and polygenic adaptations and for mapping the genetic bases of physiological and fertility phenotypes in approximately 1000 indigenous ethnically Tibetan women from Nepal, adapted to high altitude. We performed genome-wide association analysis and tests for polygenic adaptations which showed evidence of positive selection for alleles associated with more pregnancies and live births and evidence of negative selection for those associated with higher offspring mortality. Lower hemoglobin level did not show clear evidence for polygenic adaptation, despite its strong association with an *EPAS1* haplotype carrying selective sweep signals.

## Introduction

Understanding the impact of natural selection on phenotypic variation has been a central focus of evolutionary biology since its beginning as a modern scientific discipline. Decades of research have accumulated evidence for widespread adaptive phenotypic evolution in nature, including correlations between phenotypes and environmental factors [1–3], and higher reproductive success of native individuals compared to visitors [4]. Beyond the phenotypic studies, much effort has been devoted, especially in humans, to identifying adaptive alleles through indirect statistical approaches that use genetic variation data and that can detect the impact of past selective pressures [5]. The most widely used family of approaches aims at detecting new beneficial mutations that were quickly driven to high frequency or fixation by natural selection, a model that is often referred to as selective sweep [5]. These approaches have been applied to genome-wide variation data sets and have identified a large number of candidate adaptive alleles with strong effects on traits such as light skin pigmentation, lactase persistence, and resistance to pathogens [6–9]. However, genome-wide association studies have revealed that most phenotypic variation in humans is highly polygenic; in other words, it is due to the combined effects of a large number of alleles with small effects [10–12]. Under this scenario, adaptations will tend to generate upward shifts in the frequency of adaptive alleles at many loci rather than a major shift at one or few loci, as is the case, for example, for lactase persistence. Methods for detecting polygenic adaptations combine two sources of information: genome-wide association studies (GWAS) provide alleles associated with a phenotype of interest as well as their effect size, and the population frequency of GWAS alleles enable inter-population comparison [13–15]. An early example of this class of approaches showed that GWAS alleles for greater height tend to be more common in northern compared to southern European populations [13]. Indeed, height is the canonical example of polygenic adaptations in humans [14–16].

These indirect methods can provide evidence for past selective events, but each is sensitive to different selection models [17, 18], thus providing insights into a subset of adaptive alleles [19]. Moreover, these approaches cannot distinguish among selective effects on different fitness components, e.g. fertility vs. viability. A major advantage of indirect approaches is that they can detect selective sweep signals due to plausible, low selection coefficients (as long as *4N_e_s* > 1) with comparatively small sample sizes.

A complementary set of approaches aims at assessing directly the effects of genotype on reproductive fitness [20]. These direct approaches have many advantages, mainly the ability to detect selective events occurring in the present generation and the similar sensitivity to different selection models, e.g. balancing vs. directional selection [21]. However, they require large sample sizes to detect plausible selective coefficients. Large cohorts with genetic information are becoming increasingly available for humans, enabling approaches that were not feasible until recently [16, 22]. For example, a recent study analyzed two cohorts, for a total of more than 175,000 individuals, to assess genetic effects on viability by identifying alleles that changed in frequency across adulthood [23]. Another direct approach is to search for variants influencing fitness through GWAS of reproductive traits such as number of children ever born [24], twinning rate or mother’s age at first birth [25]. However, the genetic bases of reproductive traits remain markedly understudied, despite their great evolutionary and biomedical significance [26].

High altitude populations have emerged as an ideal system to study the genetic architecture of human adaptations. Populations of the high-altitude regions of Tibetan, Andean, and East African Plateaus have been exposed to the stress of hypobaric hypoxia for sufficient time [27] to have allowed the evolution of new adaptive traits [28]. In fact, these indigenous populations are known to have phenotypes distinct from those of lowlanders at high altitude and from each other. Examples include unelevated hemoglobin concentration (Hb) in Tibetans and Ethiopian Amhara [29, 30], the barrel-shaped chest in Aymara and Quechua [31] and many others (reviewed in [28, 32]). In the context of studies of high-altitude adaptation, the term “Tibetan” refers to the modern descendants of the ancient indigenous population of the Tibetan plateau who share cultural and biological affinities and reside in several polities, including Nepal. Recent population genomic studies of Tibetans detected strong selective sweep signals in Tibetans at two loci, *EGLN1* (egl-9 family hypoxia inducible factor 1) and *EPAS1* (endothelial PAS domain containing protein 1) [33–35], each coding for a key component of the regulatory program responding to variation in oxygen supply [36]. Importantly, alleles in these genes that occur at high frequency in Tibetans but are rare elsewhere were also reported to be associated with lower Hb [33–35, 37] (but see [37–39]), consistent with many observations that unelevated hemoglobin concentration is characteristic of high-altitude Tibetans.

Because the impact of hypobaric hypoxia on human physiology cannot be modified through behavioral or cultural practices, indigenous high altitude populations provide a rare opportunity to observe human evolution in action. Here, we took advantage of this property to design a study aimed at comprehesively dissecting adaptations to high altitude in a sample of ethnically Tibetan women, all citizens of Nepal who are indigenous to the country’s norther border regions, who are lifelong residents at altitudes above 3,000 m. We tested for selective events that took place in the past, through indirect approaches that can detect both selective sweeps and polygenic adaptations, as well as for ongoing events, through the direct approach of mapping measures of reproductive success. Moreover, we collected phenotype data for three physiological variables, namely total hemoglobin concentration (Hb; g/dL), percent of oxygen saturation of hemoglobin (SaO_2_) and pulse. Hemoglobin variation, in particular, is a distinctive trait in the Tibetan pattern of adaptation [32]. To gain further insights into the physiological significance of this trait, we calculated and separately mapped the concentration of oxygenated hemoglobin (“oxyHb”), which is the relevant quantity for oxygen delivery to the tissues, and of deoxygenated hemoglobin (“deoxyHb”). We found a single genome-wide significant association signal for oxyHb at the *EPAS1* locus and several signals for reproductive traits and for deoxyHb. We detected strong selective sweeps indicating past selection at *EPAS1* and *EGLN1*, while finding little evidence for polygenic adaptation toward lower Hb. We detected signatures of polygenic adaptation for reproductive traits such as numbers of livebirths and offspring mortality, consistent with selective processes that are still ongoing in contemporary populations.

## Results

### Genetic variation data of indigenous high-altitude individuals

To investigate the genetic bases of high-altitude adaptations in Tibetan populations, we collected physiological and reproductive phenotype data and saliva samples of 1,008 indigenous ethnically Tibetan women living at 3,000-4,000 m in the Mustang and Gorhka districts of Nepal (see Materials and Methods). All Tibetan participants were chosen to be 39 years of age or older so that their recorded reproductive history would have minimal confounding due to unrealized reproduction. We also obtained saliva samples for DNA extraction and analysis from103 Sherpa participants (including ten parents and offspring trios) from the high-altitude regions in the Khumbu district in Nepal. The Sherpa data were included in the reference panel for genotype imputations and in the polygenic adaptation tests, but not in GWAS (see below).

Genetic variation data of our study participants were generated by a combination of experimental and computational tools (Figure 1; also see Materials and Methods). First, we generated novel genotype data for all participants using Illumina genotyping array platforms in multiple phases (Supplementary Table 1). Briefly, all Tibetans were first genotyped for about 300K markers on the HumanCore array with additional 2,553 custom markers to cover candidate regions, including the *EGLN1*, *EPAS1*, *HIF1A* (hypoxia inducible factor 1 alpha subunit) and *NOS2* (nitric oxide synthase 2) genes. Then, we genotyped a subset of 344 unrelated Tibetans (allowing up to first cousins) and all 103 Sherpa for over 700K markers on the OmniExpress array; the same individuals were separately genotyped for two nonsynonymous SNPs in the *EGLN1* gene, rs12097901 and rs186996510 [40].

**Figure 1.**
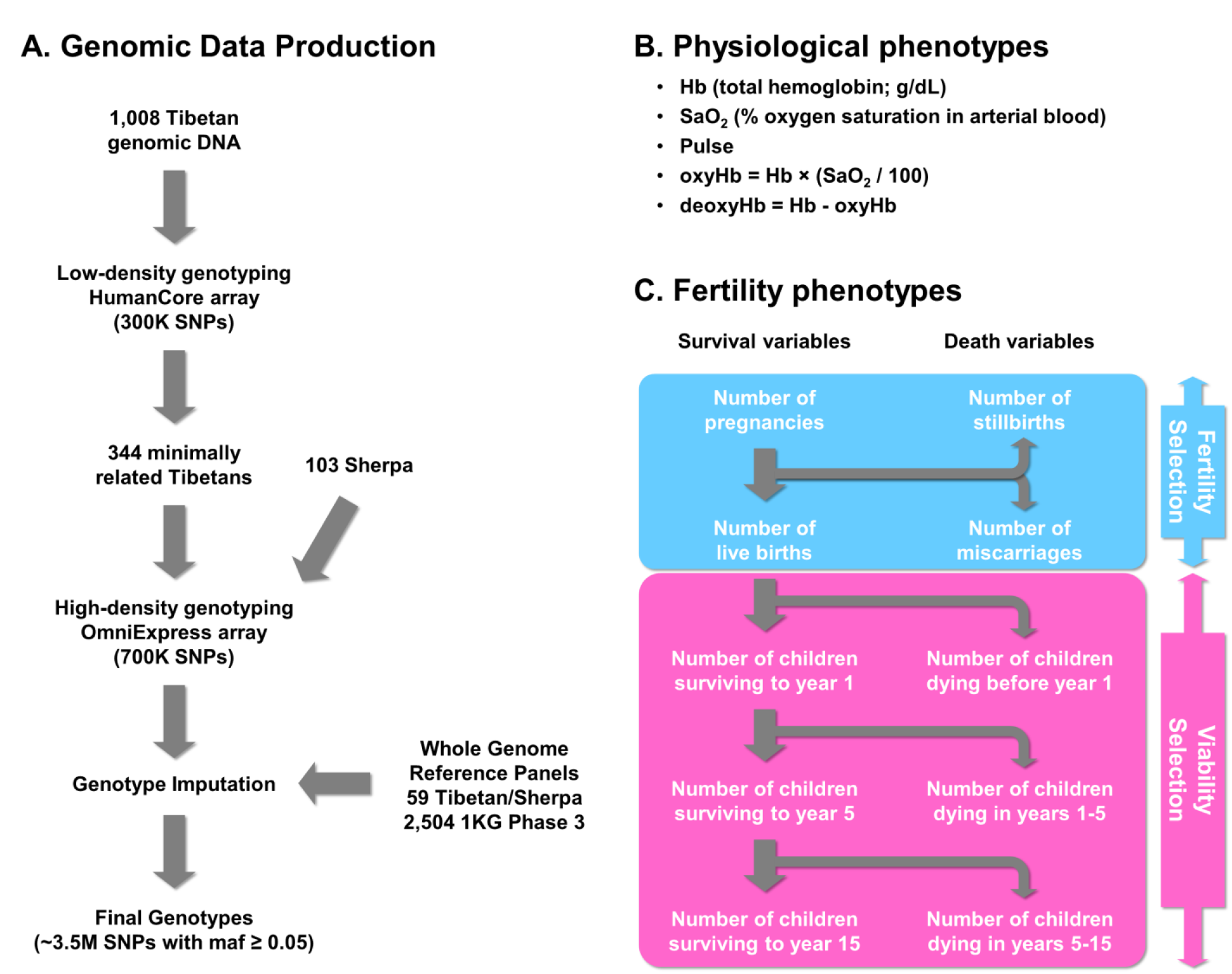
A schematic summary of the genotype and phenotype data of ethnic Tibetans in this study. (A) We array genotyped all individuals in several Illumina platforms and generated whole genome sequences for a representative subset without recent admixture. Then, all individuals went through genotype imputation using our high altitude sequence data (“high altitude panel”) and world-wide data (“1KG phase 3 panel”) as reference haplotype panels. (B) Three physiological phenotypes were directly measured in the field, and two additional ones (oxyHb and deoxyHb) were constructed from them. (C) Fertility phenotypes capture both fertility and viability selection components. For details, please see Materials and Methods.

To augment publicly available reference panels for genotype imputation, we generated whole genome sequence data of 18 Sherpa and 35 Tibetans (Supplementary Table 1). Three Sherpa trios and four Tibetan mother-daughter duos were sequenced to high coverage (~ 20x), while the remaining 36 individuals were genetically unrelated and sequenced to low coverage (~ 5x, Supplementary Table 1). For sequencing, we chose Tibetan and Sherpa individuals with no signature of recent admixture (See Materials and Methods). Adding six previously published Tibetan and Sherpa genomes [41, 42], we obtained phased genotypes of 59 individuals including 9,742,498 variants, of which 1,364,150 were not found in the 1000 Genomes Project (1KGP) phase 3 data set [43]. Among the non-1KGP variants, 540,218 were included in dbSNP150 database, while 823,932 were not. Variant annotation using the ANNOVAR program [44] identified 8,679 nonsynonymous variants, 235 nonsense coding variants, and 126 splicing variants not present in the 1KGP (Supplementary Table 2). Among the non-1KGP variants, 29.46% and 24.14% occurred as singletons and doubletons, but 11.06% of them occurred at frequency 10% or higher (Supplementary Table 2). Using both the 1KGP phase 3 data and our high altitude sequence data as reference panels, we performed genotype imputation of all samples using the IMPUTE2 program [45] to generate the analysis-ready genotype data (see Materials and Methods).

### GWAS reveals several associations with fertility phenotypes in Tibetans

We performed GWAS of 23 phenotypes characterizing the reproductive history of our study participants using a linear mixed model-based approach as implemented in GEMMA [46]. We grouped our fertility phenotypes into two categories, “fertility counts” (e.g. number of live births) and “fertility proportions” (e.g. proportion of live births among pregnancies) (Supplementary Table 3 describes the sample and summarizes the reproductive phenotypes). While the count phenotypes are more directly related to evolutionary fitness, they may be confounded by compensatory reproduction in case of a negative pregnancy outcome or by sociocultural factors that influence the count [47]. In contrast, the proportional variables are less affected by such factors and may provide information on the specific phase of the reproductive process affected by the associated genetic variation. Therefore, the counts and proportions may capture different aspects of the reproductive outcome. The GWAS was performed on the entire sample and on a subset referred to as continuously married (CM), that was composed of about 60% of participants who had been in a marital relationship throughout the ages of 25 to 40 (see Materials and Methods). This subset controls the variance in marital relationship status; on the other hand, the resulting smaller sample size reduces the power to detect significant associations.

For the fertility count phenotypes, we found 55 genome-wide significant genotype-phenotype associations clustered into five independent association signals across the genome (Table 1). First, our analysis of three fertility count phenotypes yielded genome-wide significant association peaks. Two intronic SNPs in the *CCDC141* (coiled-coil domain containing 141) gene, with the top SNP rs6711319, were associated both with the number of pregnancies (*p* = 2.10 × 10^−8^) and with the number of live births (*p* = 2.89 × 10^−9^; Figure 2 and Supplementary Figure 3). Fourteen SNPs between the *PAPOLA* (poly(A) polymerase alpha) and the *VRK1* (vaccinia related kinase 1) genes were associated with the number of stillbirths in the continuously married subset (*p* ≥ 8.38 × 10^−9^; Figure 2 and Supplementary Figure 3). The same set of SNPs also showed a suggestive association in the entire sample (*p* = 6.23 × 10^−5^ to 1.85 × 10^−4^). No expression quantitative trait loci (eQTLs) were detected in the GTEx Project data in this peak region [48], making it hard to connect the associated SNPs with a specific gene.

**Figure 2.**
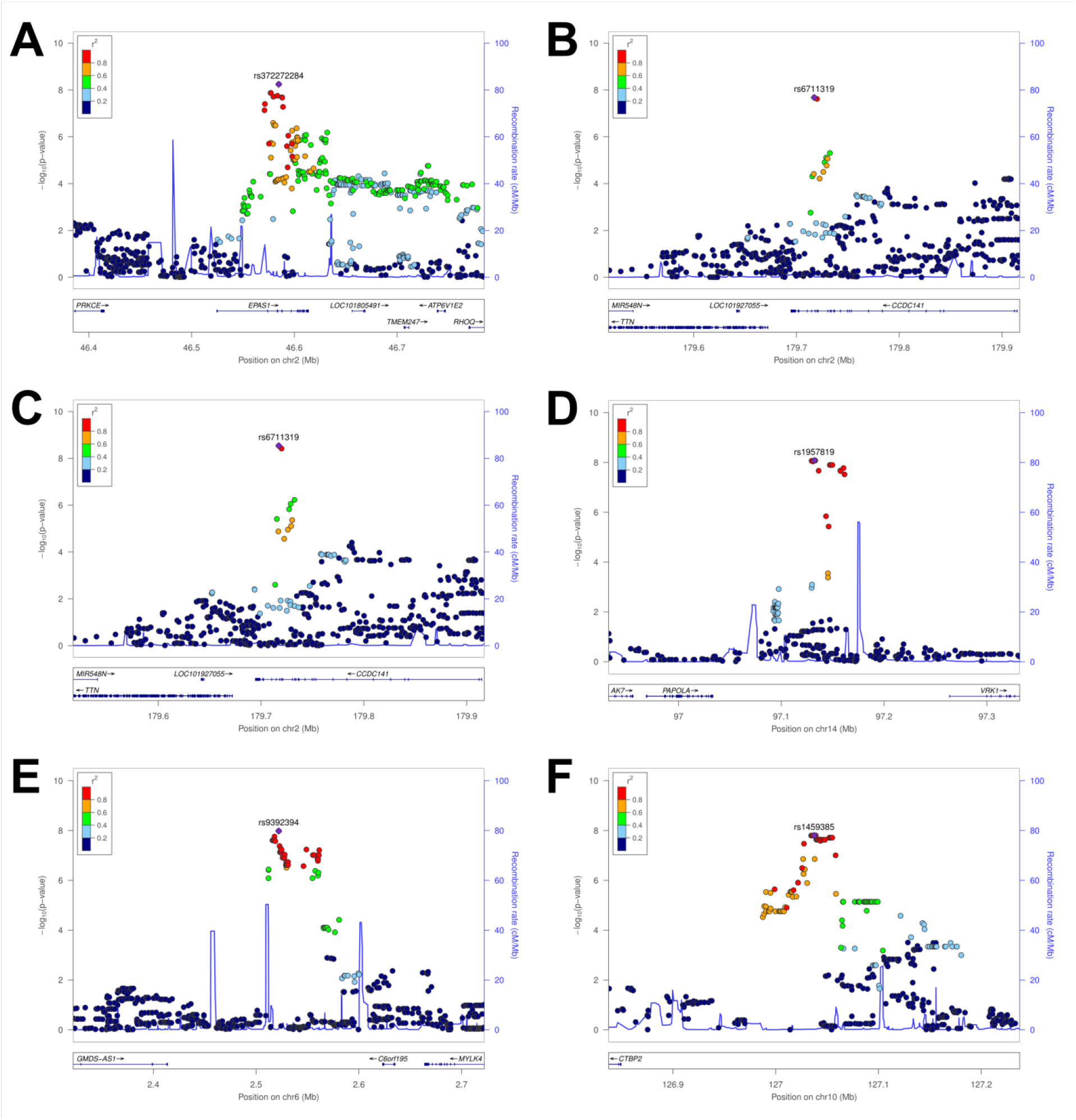
Locuszoom plots of the genome-wide significant associations found in Tibetans: (A) oxyHb and rs372272284 in the *EPAS1* gene, (B) the numbers of pregnancies or (C) live births and rs6711319 in the *CCDC141* gene, (D) the number of stillbirths and rs1957819 near the *PAPOLA* gene, (E, F) the proportion of offsprings died before age 15 years among the born alive and rs9392394/rs1459385. (A-C, E) are tests with all samples and (D, F) are those with the continuously married subset.

**Table 1.** Genome-wide significant association peaks among Tibetan women. “nSNPs” shows the number of genome-wide significant SNPs in each peak. “Top SNP” provides the rsID of the most significant SNP with effect / non-effect alleles. We chose the Tibetan minor allele as the effect allele. “Pos” is the genomic position of the top SNP in hg19 coordinates. Per allele effect size is provided in the *β* column.

| Pheno | CHR | nSNPs | Top SNP | Pos | Minor allele frequency<sup>a</sup> | $\beta^b$ | *P* | PBS | Gene |
| --- | --- | --- | --- | --- | --- | --- | --- | --- | --- |
| oxyHb | 2 | 8 | rs372272284<br>(A/G) | 46,584,859 | 0.248 | 0.386 | $5.71 \times 10^{-9}$ | 1.05 | *EPAS1*<br>(genic) |
| # of pregnancies | 2 | 2 | rs6711319<br>(G/A) | 179,717,217 | 0.248 | -0.760 | $2.10 \times 10^{-8}$ | 0.00 | *CCDC141*<br>(genic) |
| # of live births | 2 | 2 | rs6711319<br>(G/A) | 179,717,217 | 0.248 | -0.773 | $2.89 \times 10^{-9}$ | 0.00 | *CCDC141*<br>(genic) |
| # of stillbirths (CM) | 14 | 14 | rs1957819<br>(T/C) | 97,131,992 | 0.083 | 0.284 | $8.38 \times 10^{-9}$ | -0.01 | *PAPOLA*<br>(99 kb) |
| Proportion of children born<br>alive who died < 15 yr | 6 | 8 | rs9392394<br>(G/A) | 2,522,000 | 0.442 | -0.318 | $1.05 \times 10^{-8}$ | 0.00 | *C6orf195*<br>(101 kb) |
| Proportion of children born<br>alive who died < 15 yr (CM) | 10 | 29 | rs1459385<br>(T/C) | 127,037,417 | 0.089 | 0.704 | $1.59 \times 10^{-8}$ | 0.02 | *CTBP2*<br>(188 kb) |

<sup>a</sup> Minor allele frequency in our Tibetan data set
<sup>b</sup> Per minor allele effect size estimates are in the unit of g/dL (oxyHb), the number of children (the numbers of pregnancies, live births and stillbirths), or residuals in log-odds scale for fertility proportion phenotypes (proportion of children born alive who died < 15 yr)

For the fertility proportion phenotypes, two genome-wide significant association signals were detected (Table 1). Eight SNPs near *C6orf195*, with the top SNP rs9392394, were associated with the proportion of children who died before age 15 (*p* = 1.05 × 10^−8^); SNPs within 15 kb of this peak had been associated with heart, blood pressure and reticulocyte traits in GWAS [49–51]. Twenty-nine SNPs near *CTBP2*, with the top SNP rs1459385, were associated with the same phenotype in the continuously married subset (*p* = 1.59 × 10^−8^; Figure 2 and Supplementary Figure 3); other genes in this region include *TEX36* (testis expressed 36) and *EDRF1* (erythroid differentiation regulatory factor 1, which regualtes the expression of globin genes). No eQTLs were detected in the GTEx Project data in these association regions.

### EPAS1, but not EGLN1, SNPs are associated with hemoglobin levels in Tibetan women

The genetic bases of of Hb, SaO_2_, and pulse have been previously studied in outbred populations mainly of European ancestry [51–57]. Here, we performed GWAS of these key physiological phenotypes in Tibetans to potentially uncover population-specific genetic determinants. We derived two additional composite phenotypes: oxygenated hemoglobin concentration (“oxyHb”, defined as the product of Hb and SaO_2_ divided by 100), and deoxygenated hemoglobin concentration (“deoxyHb”, defined as the difference between Hb and oxyHb). Consistent with findings from other studies, these women had an average hemoglobin concentration of 13.8 g/dL ± 1.3 g/dL. Supplementary Tables 3 and 4 describe the sample and summarize the phenotypic data. Each GWAS included about 3.5 million SNPs with minor allele frequency (maf) ≥ 0.05.

Eight SNPs within a 17 kb intronic region of the *EPAS1* gene were significantly associated with oxyHb (*p* ≤ 5 × 10^−8^ for all eight SNPs, with the top signal at rs372272284; Table 1, Figure 2 and Supplementary Figure 1). Hb and oxyHb were strongly correlated (Pearson *r* = 0.874), and all eight SNPs were also strongly associated with Hb (*p* ≤ 4.10 × 10^−7^; Supplementary Table 5). This is the first report of an association of the derived Tibetan *EPAS1* alleles with a hemoglobin trait that reaches genome-wide significance levels. The results, including the estimated effect size of 0.332 g/dL per allele, support previous candidate gene studies for Hb [33, 34]. Due to strong linkage disequilibrium (LD), the signature of a selective sweep around the *EPAS1* gene in Tibetans extends farther than 100 kb; however, our large sample size and dense genetic variation data allowed us to narrow down the association signal to a 17 kb region.

Conditioning on the genotype of rs372272284, no residual association with either Hb or oxyHb was observed in the *EPAS1* locus (*p* ≥ 0.770; Supplementary Figure 2). This includes a previously identified “Tibetan-enriched” deletion (“TED”), 81kb downstream of the *EPAS1* gene, present in Tibetans but not in the introgressed Denisovan haplotype [58]. TED is in LD with the eight significant SNPs in our data set (Pearson *r* = 0.771-0.783), but its association with Hb and oxyHb was much weaker than our top SNPs (*p* = 1.64 × 10^−3^ and 3.09 × 10^−4^, respectively).

In contrast, we did not detect significant associations in the *EGLN1* gene. Two nonsynonymous variants in the *EGLN1* gene, rs12097901 and rs186996510, were previously shown to harbor strong signatures of selective sweep [37, 59]. An early study reported an association of the *EGLN1* haplotype (defined by a combination of three SNPs) with lower Hb in a sample of mixed gender with large effect size estimate 1.676 g/dL per allele [35]. However, two more recent studies, each with more than 500 Tibetan participants, reported that the derived allele of rs186996510 was associated with lower Hb only in males [37, 39]. Similarly, an analysis of *EGLN1* SNPs in 3,008 Tibetans did not detect a significant association with Hb, and reported a larger effect size for males compared to females [38]. Consistent with the more recent evidence, we did not detect a significant association at either of the two SNPs with Hb (*p* ≥ 0.224) or with oxyHb (*p* ≥ 0.268) in our Tibetan sample, which includes only females. No association was detected when we either added menopause status, a female-specific covariate of Hb, as an additional covariate or confined our analysis to post-menopausal women (*p* ≥ 0.641), thus excluding it as an explanation for the lack of association in females. We calculated that our power to detect an association with a single test (α = 0.05) for the observed allele frequency of rs186996510 (0.336) and our sample size (N = 649) was ≥ 99% for an effect size as low as 0.332 g/dL per allele, which is well below previous estimates (Supplementary Table 6). The reasons for the inconsistent findings regarding an association of *EGLN1*, an oxygen sensor in the oxygen homeostasis pathway, and hemoglobin concentration are unknown.

A recent study suggested an interaction between the effects of *EGLN1* and *EPAS1* SNPs on Hb levels [60]. However, our data showed no significant interaction between the *EPAS1* and *EGLN1* SNPs (rs372272284 and rs186996510, respectively) in the association with Hb (*p* = 0.613).

### Are lower Hb levels adaptive among Tibetan women?

The women’s reproductive history data offer a unique opportunity to ask if these selective sweep signals are associated with ongoing selection in contemporary Tibetans due to maternal factors and to estimate their selection coefficient. For each SNP, we calculated the population branch statistic (PBS), which measures the extent of allele frequency divergence on the lineage leading to a test population [34], in this case Tibetans. Consistent with previous studies, the *EPAS1* (PBS = 1.073, rs73926264) and *EGLN1* (PBS = 0.797, rs186996510) loci harbored the highest PBS values (Supplementary Table 7). Table 2 shows the association *p*-values for the top *EPAS1* Hb/oxyHb GWAS SNP and the top PBS *EGLN1* SNP with all the fertility measures included in our GWAS. There is no association between *EGLN1* and *EPAS1* SNPs and any of the direct measures of fitness, i.e. number of live births as well as number of children surviving at 1, 5 and 15 years. Nominal levels of significance are observed in some cases, but no test reaches significance after multiple test correction. Power calculations for the fertility count phenotypes suggest that we can detect such an association only if the associated selection coefficient is extremely high (≥ 6.6% per allele for 80% power given a single test; Table 2). Previous estimates of the selection coefficient for both *EPAS1* and *EGLN1*, 1.5% and 2.9%, respectively, are well below this range [37, 61]. Therefore, these results are not inconsistent with the idea that these variants are advantageous.

**Table 2.**
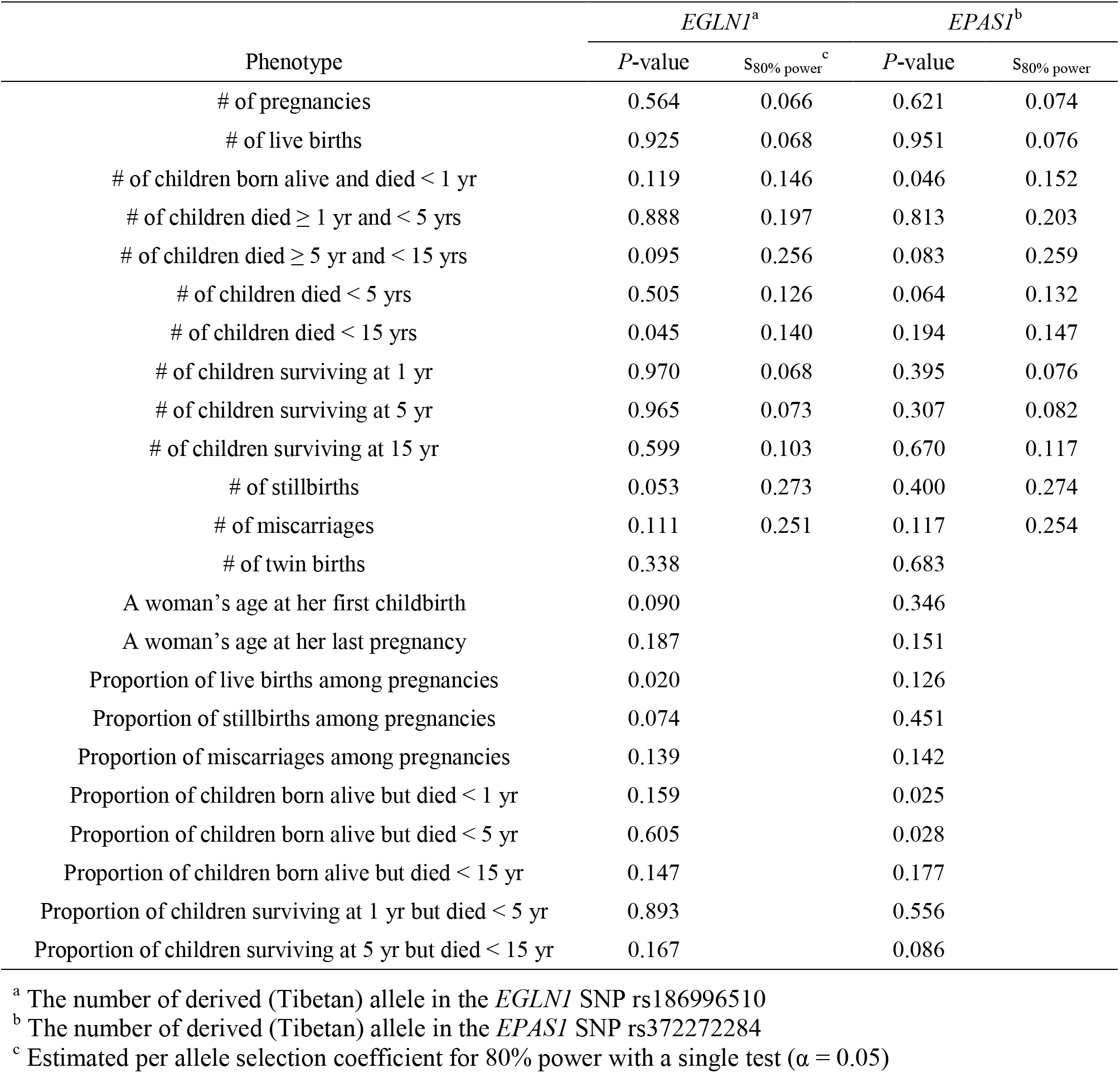
*P* values for the correlation between fertility or fertility proportion phenotypes controlling for covariates and the *EGLN1* and *EPAS1* SNP genotypes. Per-allele selection coefficient for 80% power to detect association in a single SNP test (α = 0.05) was estimated for fertility count phenotypes. No test showed *p* ≤ 0.01.

The finding of *EPAS1* alleles that are associated with lower Hb levels in Tibetans and carry a strong signature of positive natural selection led to the hypothesis that a dampening of the acclimatization response in Hb levels is adaptive in Tibetans [62]. Interestingly, *EPAS1* was the only locus among the 36 strongly associated with Hb (*p* ≤ 10^−4^) to carry a strong signature of selective sweep. Importantly, although the *EPAS1* SNPs are the only genome-wide significant association signal, they explain only 2.7% of the inter-individual variation in Hb suggesting that many other loci with effects on Hb exist in Tibetans. Consistent with the idea of polygenic inheritance for Hb, a recent GWAS revealed 140 loci with small effects contributing to Hb levels in a large sample of European ancestry [51]. Therefore, if lower Hb were adaptive, we would expect to observe signatures of polygenic adaptation at Hb-associated SNPs as a group, namely a trend towards lower frequencies of the Hb-increasing alleles at many unlinked SNPs in high altitude compared to low altitude samples. To test for this possibility, we used two methods specifically designed to detect consistent changes in the frequency of alleles at many unlinked Hb GWAS SNPs.

First, we calculated the mean allele frequency difference of Hb-increasing alleles between Tibetans or Sherpa and 1KGP CHB (Han Chinese in Beijing, China) across the 36 and 43 independent SNPs (*p* ≤ 10^−4^) ascertained from our Tibetan Hb and oxyHb GWAS, respectively. Compared to 10,000 sets of control SNPs [13], the Hb SNPs identified in Tibetans showed on average a lower frequency of Hb-increasing alleles in both Tibetans and Sherpa, suggesting selection favoring lower Hb levels (one-sided empirical-p = 0.047 and 0.018, respectively; Figure 3). The Tibetan oxyHb SNP set also showed a similar pattern (*p* = 0.102 and 0.046 for Tibetans and Sherpa, respectively; Supplementary Figure 4). However, when the *EPAS1* SNP rs372272284 was excluded, no difference between the Hb- or oxyHb-associated SNPs and control SNPs was observed (*p* ≥ 0.211; Figure 3 and Supplementary Figure 4). Thus, the overall frequency difference seemed entirely due to the large frequency differentiation of the *EPAS1* SNP: the Hb-increasing allele frequency was 0.253 and 0.990 for Tibetans and CHB, respectively.

**Figure 3.**
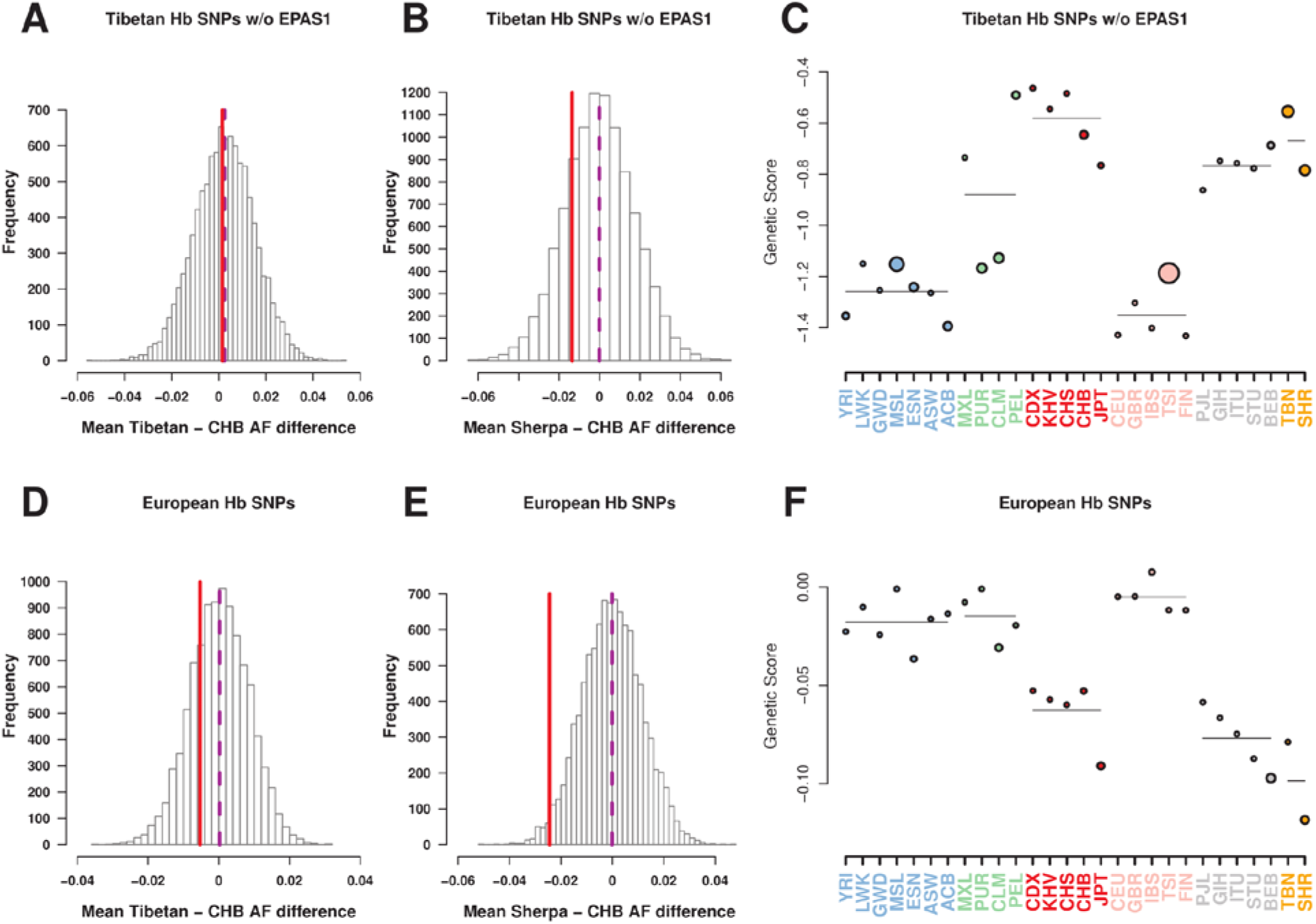
Tests of polygenic adaptation of Hb-associated SNPs: (A-C) 35 SNPs from our Tibetan GWAS (*p* ≤ 10^−4^) after excluding the *EPAS1* SNP rs372272284, and (D-F) 96 genome-wide significant SNPs from a large GWAS of mostly European cohorts. The mean frequency differences of trait-increasing alleles between Tibetans and CHB (A, D) and between Sherpa and CHB (B, E) were presented (solid red line) together with the empirical null distribution of 10,000 sets of matched random SNPs. (C, F) The genetic values of populations (filled dots) and of regions (horizontal lines) were plotted. The size of dots and the width of lines are proportional to the significance of the corresponding outlier test.

**Figure 4.**
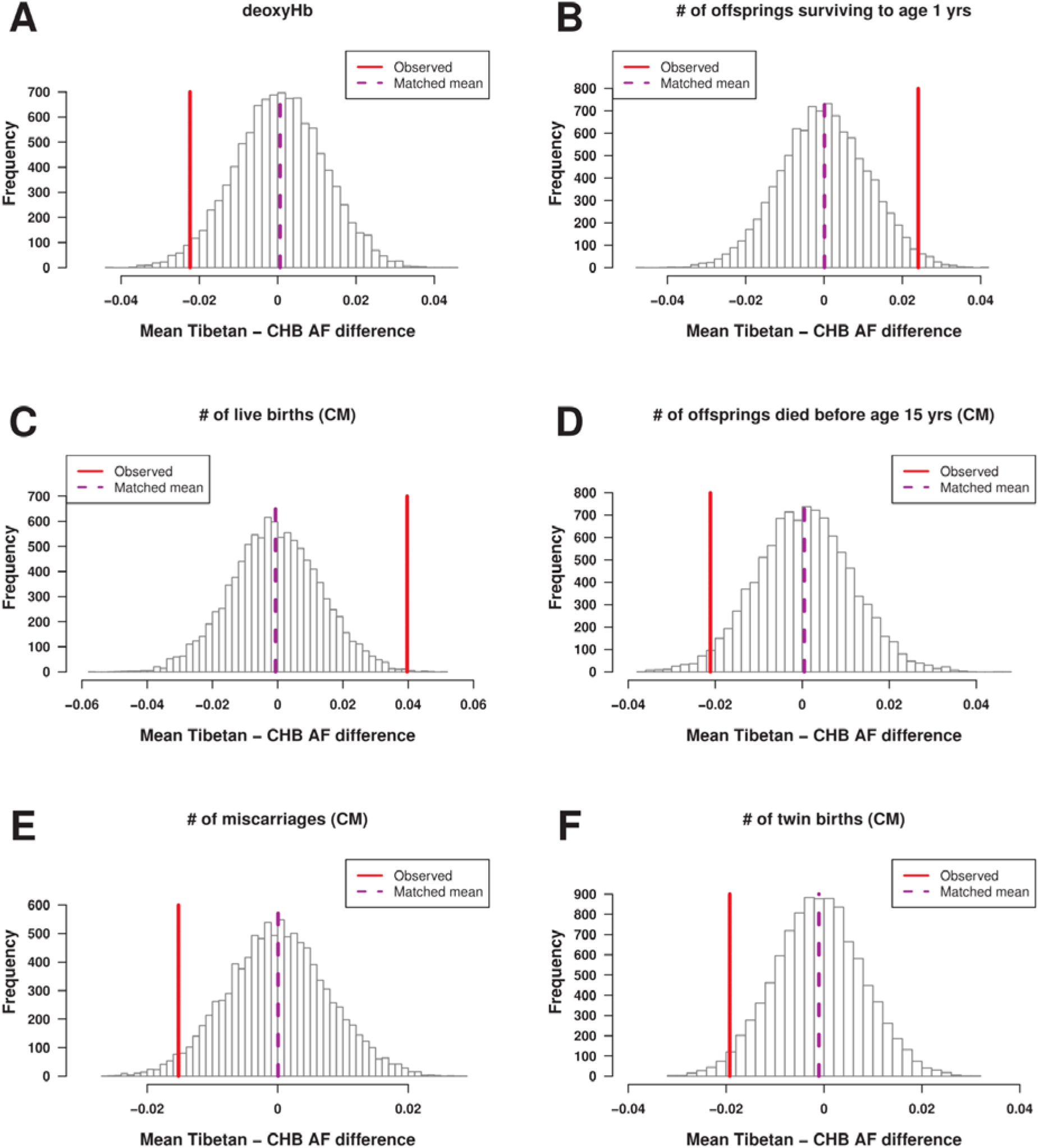
Phenotypes showing signatures of polygenic adaptations in Tibetans. The mean frequency difference of trait-increasing alleles was presented (solid red line) together with the empirical null distribution of 10,000 sets of matched random SNPs. (C-F) uses GWAS SNPs from the “CM” subset.

GWAS SNP effect sizes have been shown to be correlated between European and East Asian populations [63], implying that SNPs identified in the large Hb GWAS in Europeans may be informative about the genetic bases of Hb variation in our Tibetan sample. Therefore, we also tested for polygenic adaptation using SNPs identified in a large European GWAS [51]; we used the 94 GWAS SNPs that were called in our data set. This set of SNPs did not include any *EPAS1* SNPs, because the Tibetan *EPAS1* haplotype is virtually absent outside Tibetan populations [64]. We found a trend towards lower frequencies of Hb-increasing alleles in both Tibetan and Sherpa, but this trend was significant only in Sherpa (*p* = 0.249 and 0.019 for Tibetans and Sherpa, respectively; Figure 3).

We also applied a more recently developed set of tests for polygenic adaptation [14], using our data from Tibetans and Sherpa along with the 1KGP populations. This approach calculates a genetic value for each trait in each population by summing up the product of the frequency at each GWAS SNP and the effect size of that SNP and it compares GWAS-ascertained SNPs with a large number of control SNPs. Specifically, we focused on two tests. The “overdispersion” test asks if allele frequencies of the GWAS SNPs as a group show either unusually big differentiation across populations or unexpectedly strong correlation in the direction of change. The “outlier” test asks if the genetic value of a trait in a population or a group of populations is significantly different from that of the other populations. Excluding the *EPAS1* SNP, neither the overdispersion test nor the outlier test for the high-altitude populations yielded results reaching nominal levels of significance (*p*_overdispersion_ = 0.695 and *p*_outlier_ = 0.066 with no multiple test correction) for oxyHb, or Hb (*p*_overdispersion_ = 0.201 and *p*_outlier_ = 0.846 with no multiple test correction) (Figure 3, Supplementary Figure 4 and Supplementary Table 8). In contrast (but consistent with the pairwise population test above), the outlier test was highly significant when the *EPAS1* SNP was included (*p* ≤ 0.0008; Figure 3, Supplementary Figure 4 and Supplementary Table 8). Using the Hb associated SNPs from the European GWAS, we again observed a trend toward lower genetic values in the high altitude populations, but it did not reach levels of statistical significance (*p*_overdispersion_ = 0.433 and *p*_outlier_ = 0.110 with no multiple test correction). Therefore, these analyses do not provide evidence that alleles associated with lower Hb levels were selected for, except for the *EPAS1* locus. Given that the *EPAS1* SNPs explain a small fraction of the total variation in Hb levels, these results raise the question of whether unelevated Hb *per se* was the adaptive trait in Tibetans.

Among the other physiological phenotypes, deoxyHb alone showed polygenic adaptation signals, as both the outlier and the pairwise difference tests were significant (*p*_outlier_ = 0.023 and *p*_pairwise_ = 0.002; Supplementary Figure 4 and Supplementary Table 8). This result is not inconsistent with the lack of evidence for polygenic adaptation toward lower Hb because this is not strongly correlated with deoxyHb (r = 0.441 compared to that with oxyHb *r* = 0.874). The alleles associated with higher deoxyHb in Tibetans were on average less common in Tibetans than in 1KGP CHB and the genetic values of deoxyHb in Tibetans and the Sherpa tended to be lower than those in 1KGP East Asians. Maximizing oxygen delivery while minimizing blood viscosity is likely to be beneficial in high-altitude environments; therefore, this advantage may underlie our signal of polygenic adaptation for lower deoxyHb. Neither SaO_2_ nor pulse provided a significant result for any polygenic adaptation test (Supplementary Table 8).

### Multiple reproductive fitness traits show evidence of polygenic adaptation in Tibetans

The GWAS of reproductive traits allowed us to identify candidate variants that are currently being selected for in the sampled Tibetan population. In our results, none of the most strongly associated variants with reproductive outcomes showed strong signals of selective sweeps. However, if these variants indeed affect reproductive fitness, we might expect signals of polygenic adaptation. To test this idea, we used the same tests described above.

Interestingly, a number of reproductive traits showed strong signatures of polygenic adaptations based on the outlier test; the pairwise population difference test, which uses less information and hence is likely to be less powerful, gives broadly consistent results, although at lower levels of significance (Table 3, Supplementary Figure 5 and Supplementary Table 8). Contrary to the case of Hb, we see significant polygenic adaptation signals in several measures directly related to reproductive fitness. Interestingly, the significant signals are observed for both the viability (e.g. the number of children born alive but died before 15 years; *p*_outlier_ = 0.000, *p*_pairwise_ = 0.024) and the fertility fitness component (e.g. the number of live births; *p*_outlier_ = 0.002, *p*_pairwise_ = 0.002). Furthermore, consistent with expectations, alleles increasing offspring mortality were selected against whereas those increasing offspring survival were positively selected for. A variable known to be directly linked to reproductive fitness, i.e. a woman’s age at her first childbirth, is also under selection, with earlier ages being advantageous, as expected. Twinning appears to have been selected against in Tibetan women. Although twinning may increase fitness, it is also associated with increased risks to mother and offspring due to limits on women’s ability to support adequate weight gain for two babies during the third trimester and to the lower birth weight of twins relative to singletons [65], which in turn is associated with higher neonatal and infant mortality.

**Table 3.** Results of polygenic adaptation tests for a chosen set of physiology and fertility phenotypes. Results for all GWAS phenotypes are presented in Supplementary Table 8.

| Phenotype | Direction<sup>a</sup> | GWAS sample<sup>b</sup> | Outlier test | | $P_{\text{frq.diff}}^{\text{c}}$ |
| --- | --- | --- | --- | --- | --- |
| | | | Statistic | $P$-value | |
| 1. Fertility phenotypes |  |  |  |  |  |
| A woman’s age at her first childbirth | - | CM | -1.9286 | 0.0000 | 0.576 |
| # of live births | + | CM | 2.5062 | 0.0020 | 0.002 |
| # of miscarriages | - | CM | -3.0936 | 0.0000 | 0.024 |
| # of children born alive but died < 1 yr | - | CM | -2.4325 | 0.0000 | 0.012 |
| # of children born alive but died < 15 yr | - | CM | -2.0293 | 0.0000 | 0.024 |
| # of children born alive but died < 5 yr | - | All | -1.5043 | 0.0000 | 0.134 |
| # of children died ≥ 5 yr and < 15 yrs | - | CM | -1.1255 | 0.0256 | 0.304 |
| # of children surviving at 1 yr | + | All | 1.7603 | 0.0100 | 0.017 |
| Proportion of children born alive but died < 15 yr | - | CM | -1.2639 | 0.0224 | 0.101 |
| Proportion of stillbirths among pregnancies | - | CM | -2.1763 | 0.0008 | 0.230 |
| Proportion of stillbirths among pregnancies | - | All | -1.6320 | 0.0004 | 0.285 |
| # of twin births | - | CM | -2.4344 | 0.0000 | 0.021 |
| # of twin births | - | All | -1.1421 | 0.0024 | 0.139 |
| 2. Physiological phenotypes |  |  |  |  |  |
| Hb | - |  | -1.0490 | 0.0000 | 0.047 |
| Hb without *EPAS1* | - |  | -0.5003 | 0.8460 | 0.467 |
| oxyHb | - |  | -0.5657 | 0.0008 | 0.102 |
| oxyHb without *EPAS1* | - |  | 0.8596 | 0.0656 | 0.554 |
| deoxyHb | - |  | -1.4628 | 0.0024 | 0.023 |
| Pulse | - |  | -0.4412 | 0.4048 | 0.062 |

<sup>a</sup> Assumed direction of positive selection for each phenotype
<sup>b</sup> GWAS sample sets for fertility phenotypes: all samples (All) and continuously married subset (CM)
<sup>c</sup> One-sided empirical *p*-value for the mean frequency difference test

### Hb and pulse are correlated with aspects of reproductive success in Tibetan

A previous analysis of this sample of Tibetan women found strong relationships between physiological traits and reproductive success in this Tibetan sample by using a large set of covariates, including physiological, sociocultural, and socioeconomic variables (e.g. relative wealth rank, type of marriage, and marital status) [47]. Because the present analyses used only a subset of covariates used in the previous study, we tested for association of physiological traits and reproductive success by correcting for the same set of covariates used in our GWAS. Consistent with the previous analysis, we found that lower Hb correlated with a higher proportion of live births among pregnancies (*p* = 0.002). We also found that Hb correlated positively with the numbers of stillbirths or miscarriages (*p* = 0.040 and 0.057, respectively), as well as their proportions among pregnancies (*p* = 0.023 and 0.033 for stillbirths and miscarriages, respectively).

Another interesting finding was the negative correlations between pulse and most of the fertility phenotypes, with the strongest correlations found with the numbers of pregnancies and livebirths (*p* = 2.02×10^−5^ and 2.76×10^−5^, respectively; Supplementary Table 9). Pulse’s negative association with a woman’s age at her last pregnancy partially accounts for this strong correlation; however, the association between pulse and the number of pregnancies remained significant (*p* = 0.005) after correcting for a woman’s age at her last pregnancy, even if weaker. The pulse and fertility traits were previously shown to be marginally correlated if a larger set of covariates was included in the model (*p* = 0.130 and 0.069 for the numbers of pregnancies and livebirths, respectively) [47].

## Discussion

We identified several genome-wide significant associations with key physiological and fertility phenotypes in Tibetans (Figure 2 and Table 1), by analyzing new dense genome-wide variation data of over 1,000 indigenous inhabitants above 3,000 m in Nepal (Supplementary Table 1). Using genetic variants identified in our GWAS, we found that several phenotypes showed signatures of polygenic adaptation towards better reproductive outcomes (e.g. the number of livebirths) (Table 3, Supplementary Figure 5 and Supplementary Table 8). Surprisingly, we did not find clear evidence for polygenic adaptation towards low Hb in Tibetans beyond a link through the *EPAS1* gene, even though we confirmed a correlation between low Hb and better reproductive outcomes. Because Hb concentration is a polygenic trait, these results raise the question of whether lower hemoglobin is causally related to higher reproductive fitness.

Our GWAS of fertility phenotypes discovered three genome-wide significant associations (Table 1 and Supplementary Figure 3). Those signals lie in or near genes of potential biological relevance. First, the association peak for the number of pregnancies and of livebirths is located within an intron of the *CCDC141* gene (Figure 2), which is expressed in the heart and had been linked to a rare form of hypogonadotropic hypogonadism [66]. This gene is an immediate neighbor of the *TTN* (titin) gene, which codes for a major component of cardiac muscle and has been linked to idiopathic dilated and peripartum cardiomyopathy and cardiac remodeling [67, 68]. Genetic variants within 6 kb from our association peak were reported to be associated with cardiac phenotypes, such as heart rate [52, 69]. Although these GWAS signals were not associated with pulse, we hypothesize that they influence heart function, which in turn may affect pregnancy outcomes in the extreme high-altitude environments. The observed negative correlation between pulse and the number of livebirths is consistent with this idea.

Second, the top SNP in chromosome 14 associated with the number of stillbirths is 99 kb away from the *PAPOLA* gene encoding a poly-A tail polymerase that affects mRNA stability and nuclear export. Intriguingly, the product of this gene is inhibited by cordycepin, an adenosine analog (3’ deoxyadenosine), found in nature in a fungus, “Yartsa gunbu” or *Cordyceps sinensis*, which is native to the highlands of Nepal and Tibet. Harvest of this fungus for sale primarily in China is a major source of household revenue in the Gorkha district, from where about one third of our participants were recruited. Although it is not a species consumed by ethnic Tibetan women in this region, our results raise the possibility that the *PAPOLA* SNPs may affect the stillbirth phenotype by interacting with an exposure to *C. sinensis* during pregnancy. An alternative and equally likely explanation is that these SNPs influence reproductive outcomes through mechanisms not involving cordycepin exposure, for example by affecting mRNA levels of key genes involved in inflammatory processes, as suggested in knockdown experiments of the *PAPOLA* gene [70], or through mechanisms involving other nearby genes.

The availability of reproductive history data in a population with little or no birth control offers unique opportunities for elucidating the adaptation process. Indeed, the ethnic Tibetan women sampled in this study have high birth rates (the number of livebirths = 5.38 ± 2.79; mean ± 1 standard deviation) and live in a mostly traditional society, where modern medical care, including in some regions contraception, has been introduced only very recently [47]. The reproductive data allowed testing for a relationship between genetic or phenotypic variation and fitness differential. Interestingly, genetic variation carrying well-established signals of selective sweeps, i.e. *EGLN1* and *EPAS1* SNPs, was not associated with reproductive success (Table 2 and Supplementary Table 9). A likely explanation for these observations is that we do not have power to detect an association: we estimate that only very strong positive selection (per-allele selection coefficient ≥ 6.6%) can be detected given our sample size (Table 2). However, we did detect significant signals of polygenic adaptations using the SNPs identified in our GWAS of fertility variables. Importantly, alleles increasing survival variables were selected for while those increasing death variables were selected against, as expected (Table 3 and Supplementary Table 8). These findings connect a selection signal identified by using a population genetics approach and measures of reproductive success and point to ongoing natural selection.

An attenuated erythropoietin and Hb concentration response to hypobaric hypoxia is a hallmark phenotype of the “Tibetan pattern” of high-altitude adaptations, which is markedly different from that of Andean highlanders [32, 71, 72]. The low prevalence among Tibetans of diseases associated with elevated Hb concentration, such as chronic mountain sickness [73], and a signal of selective sweep in the *EPAS1* gene [33, 34] have led to the hypothesis that unelevated Hb is adaptive in Tibetan highlanders [62]; this hypothesis was also substantiated by the correlation between low Hb and better reproductive outcomes in our Tibetan sample [47]. Our GWAS provides the first genome-level support for the association between the Tibetan *EPAS1* haplotype and low oxyHb, which correlates highly with total Hb. Interestingly, the association was stronger for oxyHb than for total Hb (Table 1 and Supplementary Table 5), while it was not significant for deoxyHb (*p* = 0.883 for rs372272284). This observation raises the possibility that it is the oxygen-carrying portion of total Hb that drives the well-replicated association between *EPAS1* SNPs and Hb. We also found that SNPs associated with Hb, either in our own GWAS in Tibetans or in a much larger one in Europeans [51], did not show polygenic adaptation signals in our Tibetan sample, if the *EPAS1* SNP was excluded from the analysis (Figure 3). Intriguingly, the Sherpa, who are closely related to other Tibetan populations and also have unelevated Hb levels [41, 71, 74], show a significant trend towards lower frequencies of the Hb-increasing alleles in one of the two polygenic adaptation tests (*p* = 0.019), despite the smaller sample size compared to our Tibetans. Based on our estimate of 0.386 g/dL per allele, and a mean allele frequency difference of 0.743 between high and lowlanders, we calculated that the *EPAS1* SNPs can explain 52% of the 1.1 g/dL difference reported in [75] between Tibetan and Han Chinese women in the same age range. In our sample, the *EPAS1* SNP explain only 2.7% of inter-individual variation in Hb: therefore, almost all within-population (97.3%) as well as a substantial portion of between-population (48%) variation remains unexplained.

Several scenarios could account for these results. Incomplete power in the Tibetan GWAS and/or in the polygenic adaptation tests could underlie the lack of clear evidence for polygenic adaptation for lower Hb levels, although we had sufficient power to detect polygenic adaptation signals for several other traits in the same samples. The lack of evidence supporting low Hb as the selected trait in Tibetans stands in stark contrast with the strong selective sweep signal at *EPAS1* and with the significant evidence for polygenic adaptations toward lower deoxyHb. This finding raises the question of whether unelevated Hb was the true target of selection in Tibetans rather than a mere correlate of the true adaptive trait. This scenario would be consistent with the observed correlation between low Hb and better reproductive outcomes because pleiotropy can induce a non-causal association between phenotypes. A recent study showed that the same *EPAS1* SNP that is associated with Hb and other hematological traits is also associated with uric acid levels [38], suggesting that indeed SNPs in *EPAS1*, a transcription factor with dozens of target genes, may affect multiple, seemingly unrelated phenotypes. Interestingly, the peak of our association signal for oxyHb at *EPAS1* spans active enhancer (H3K27Ac) marks in human umbilical endothelial cells, as detected by the ENCODE project [76]. Therefore, it could be speculated that the SNPs that influence variation in oxyHb/Hb levels also affect endothelial function and vascularization, with beneficial effects in oxygen delivery at high altitudes. These findings suggest that the WHO altitude-adjusted elevated hemoglobin cut-off for detecting iron-deficiency anemia [77] may be inappropriate for use among Tibetan women, a result of this work that has public health implications and that warrants further research.

This study was designed to extend the genetic study of human local adaptation beyond selective sweeps and candidate gene associations, by collecting genotype and physiological phenotype and reproductive history data for a large group of indigenous high-altitude Tibetan women in Nepal. Using this data set, we successfully identified several new genome-wide associations and signatures of polygenic adaptations. Our sample size of 1,000 participants is remarkably large for the genetic study of populations living in remote locations in a traditional society, but we acknowledge that is rather small for a modern-day GWAS. The census population size of ethnic Tibetans of villages in this region set the ultimate constraint on our sample size, which was obtained by recruiting virtually all inhabitants fitting our inclusion criteria. Despite this constraint, this study shows the necessity to study phenotypes of locally adapted populations in their native environments to correctly identify the adaptive phenotypes. With ever increasing throughput to generate genetic and phenotypic variation data, in-depth phenotyping of potentially adaptive features will help better understand how Tibetans and other populations living in extreme environments have adapted to their habitats.

## Materials and Methods

### Sample information

A total of 1,008 ethnic Tibetan participants were recruited from high-altitude villages in Mustang and Ghorka districts in Nepal in the spring and summer of 2012. All participants were women of age 39 or older and lifelong residents above 3000 m of altitude. The study communities in Nepal lie on the southern aspects of the Tibetan Plateau. Although they are citizens of Nepal, local people speak Tibetan dialects, practice forms of religion and social organization akin to those across the Tibetan Plateau, and retain the characteristic agro-pastoral and trading mode of subsistence common among highland Tibetans [47]. An additional 103 Sherpa participants were recruited from high-altitude villages in the Khumbu district in Nepal in the summer of 2014. Most of the Sherpa participants were women of age 39 or older. We collected saliva samples of husbands and children for 12 of them. Saliva samples were collected in the field using OG-500 Oragene DNA collection kits (DNA Genotek Inc., Otawa, ON, Canada) and genomic DNA (gDNA) were extracted using the prepIT-L2P reagents (DNA Genotek Inc) following the manufacturer’s protocol. Blood hemoglobin concentration (Hb), percent arterial blood oxygen saturation (SaO_2_), and pulse rate (pulse/minute) were measured as described in Cho et al. [47]. Two additional phenotypes, oxygenated and deoxygenated hemoglobin concentrations (oxyHb and deoxyHb, respectively), were calculated from Hb and SaO_2_ as follows: oxyHb = Hb × SaO_2_ / 100 and deoxyHb = Hb – oxyHb. For each participant, an interview session was held to retrieve detailed reproductive history as well as to collect other potential covariates. The study protocol was approved by the institutional review boards at Case Western Reserve University, Dartmouth College, Washington University at St. Louis, by the Nepal Health Research Council and by the Oxford Tropical Research Centre Ethics Committee. A written informed consent was signed by each participant. A summary of the Tibetan samples and their phenotype measurements are presented in Supplementary Table 4. Detailed description of the Tibetan samples, the phenotype and covariate data collection was published in Cho et al. [47].

### Array genotyping

We generated new genome-wide genotype data for a total of 1,134 individuals indigenous to the high-altitude regions in the Himalayas in Nepal, including 1,001 ethnic Tibetans from the present study and 103 Sherpa (Supplementary Table 1). Array genotyping was performed in two phases. First, all Tibetan individuals were genotyped on 301,299 biallelic markers using the customized Illumina HumanCore-12 v1.0A array, which includes probes for additional 2,553 markers from 19 genomic loci presumed adaptive in Tibetans including the *EPAS1*, *EGLN1*, *HIF1A* and *NOS2* genes. Then, a subset of 344 unrelated Tibetans from the present study and all 103 Sherpa individuals were genotyped on 716,503 markers using the Illumina OmniExpress-24 v1.0 array to obtain denser genome-wide variation data. For each array platform, genotypes were called using the genotyping module in the Illumina Genome Studio with default parameters (GenCall score threshold 0.15). Previously defined clusters, downloadable from the Illumina website, were applied for genotype calling. For the 2,553 custom markers we added to the HumanCore array, we retrieved intensity data from the Illumina Genome Studio and performed genotype calling using the OptiCall v0.6.4 [78]. For 344 Tibetans genotyped on both Illumina platforms, we used genotype calls from the HumanCore array for the overlapping 253K markers. Genotype calls from the two platforms were highly concordant, with the average 99.98% concordance.

### Genotyping of nonsynonymous EGLN1 SNPs

We separately genotyped two non-synonymous SNPs in the *EGLN1* gene, rs12097901 and rs188966510, in the set of 344 unrelated Tibetans. We used Epicenter FailSafe^TM^ PCR system with the manufacturer’s recommended condition in buffers G and H, instead of using standard TAQ polymerases. We generated a 1,025 bp PCR fragment in an 11 ul reaction volume using a previously published primer pair PHD2-X1F (CCCCTATCTCTCTCCCCG) and PHD2-X1R (CCTGTCCAGCACAAACCC) [59]. These PCR products were sequenced using BigDye^®^ Terminator v3.1 cycle sequencing kit and the PHD2-X1F primer in an Applied Biosystems^TM^ 3730XL DNA Analyzer. In a few cases where initial amplification failed, samples were diluted 4x in water, which in most cases allowed successful subsequent amplification. Genotypes were scored manually from chromatograms.

### Sample selection for whole genome sequencing

We generated novel whole genome sequence data for 18 Sherpa and 35 Tibetans from the present study, all from Nepal. Seventeen individuals were sampled with known familial relationships (four Tibetans mother-daughter duos and three Sherpa parents-offspring trios), and sequenced to high-coverage (around 20x autosomal coverages) to generate high quality phased genome sequences. The remaining 36 individuals were chosen to be unrelated and sequenced to low-coverage targeting 5x autosomal coverage.

For Sherpa, we began with 172 individuals, including 103 newly genotyped in this study and 69 previously published [41], and chose a subset of 101 unrelated individuals allowing first cousins. Coefficients of relatedness were calculated using PLINK v1.07 [79]. Then, we estimated population structure in these unrelated Sherpa, together with 30 Tibetans from near Lhasa [80] and 103 1KGP CHB, using an unsupervised genetic clustering algorithm in ADMIXTURE v1.22 [81]. Using estimates from K=2, we chose 51 Sherpa with > 95% of their ancestry from a component enriched in Sherpa and Tibetans (the remaining portion come from an ancestry representing CHB-related low altitude East Asians). Among them, we chose three pairs of couples with their offspring and 9 additional unrelated individuals for high- and low-coverage sequencing, respectively.

For Tibetans, we ran ADMIXTURE with K=3 in a supervised mode, with 103 1KGP CHB, 103 1KGP GIH (Gujarati Indians in Houston, Texas) and the 51 unrelated Sherpa as three reference groups. Pairwise relatedness was then calculated with the ADMIXTURE output using the RelateAdmix v0.08, controlling for population structure due to admixture [82]. Among individuals with minimum South Asian ancestry (< 1%), represented by GIH, we chose four pairs of mother-daughter duos of Tibetans from the present study and 27 unrelated individuals for high- and low-coverage sequencing, respectively.

### Whole genome sequencing

Single-barcoded libraries for Illumina sequencing were constructed using the TruSeq library preparation kit. Libraries were pooled into multiple batches and sequenced in the Illumina HiSeq 2500 and 4000 machines for paired-end (PE) 100 and 125 bp designs (Supplementary Table 1). Sequence reads were demultiplexed with no mismatch in 6-bp barcode sequence allowed. Reads were mapped to the human reference genome sequences (hg19) downloaded from http://hgdownload.soe.ucsc.edu/goldenPath/hg19/chromosomes/, using BWA backtrack v0.7.4 with-q15 option [83]. PCR and optical duplicate reads were marked using Picard tools v1.98 (http://broadinstitute.github.io/picard/) and were excluded from further analysis. Local realignment around indels and base quality score recalibration were performed using the GenomeAnalysis ToolKit (GATK) v2.8-1, following the best practice pipeline [84–86]. Finally, analysis-ready BAM files for variant discovery and genotype calling were produced using Samtools v1.2 [87] by filtering out reads with Phred-scaled mapping quality lower than 30.

LD-aware variant and genotype calling was performed using the GotCloud pipeline [88] with default parameters. The analysis-ready BAM files of 53 newly sequenced individuals and 6 previously reported ones, four Sherpa and two Nepali Tibetans [41, 42], were provided to the pipeline together.

### Imputation of array genotype data

We performed genotype imputation of Tibetan and Sherpa samples, which were array-genotyped either in the present or in our previous study [41] (Supplementary Table 1). For each array genotyping platform, low quality markers and samples were filtered out by applying the following filters: per-marker missing rate ≤ 0.05, Hardy-Weinberg equilibrium (HWE) *p*-value ≥ 0.00001 and per-individual missing rate ≤ 0.03. Strand-ambiguous (A/T and G/C) SNPs were removed and only SNPs in autosomes or X chromosome were retained for imputation. The filtering process was performed using PLINK v1.90 [89]. Genotype imputation was performed for each set of samples separately using IMPUTE2 v2.3.2 [45]. We used both our phased genotype calls of 59 high-altitude samples and the 1KGP phase 3 reference data set, downloadable from <u>https://mathgen.stats.ox.ac.uk/impute/1000GP_Phase3.html</u>, as imputation references by merging them with “-merge_ref_panels” flag in IMPUTE2. For other parameters, we used default values set by the program. Following imputation, genotypes with posterior probability ≥ 0.9 were accepted. Genotypes were assumed to be missing if none of three possible genotypes reached posterior probability threshold of 0.9. Then, we conducted an additional round of quality control by removing SNPs with missing rate higher than 0.05 or HWE p-value smaller than 10^−6^.

### Genome-wide association analysis

Among 1,001 successfully genotyped and imputed Tibetan women, 991 individuals were included in our genome-wide association analysis (GWAS). Four individuals were excluded from the analysis because they were born below 3,000 m). Another individual was excluded from the analysis was a genetic outlier who clustered with individuals from the Indian subcontinent. The other five were excluded either because they had inconsistent reproductive record or because they were nuns who became celibate during their reproductive years.

For physiological phenotypes, we chose relevant covariates by performing a stepwise model selection, allowing removal of a single covariate each step if likelihood ratio test (LRT) p-value obtained from the “lrtest” function in the R “lmtest” library was bigger than 0.05. The final sets of chosen covariates for physiological covariates are listed in Supplementary Table 3. For fertility phenotypes, we used an *a priori* chosen set of four covariates: age, subdistrict, use of contraception and “continuously married (CM)” status. Use of contraception was categorized into three classes: never used, previously used, and currently in use. “Continuously married” status is a binary variable defined as being in a marital relationship throughout the ages of 25 and 40. It includes two who had experienced less than two years of gap before re-marriage following divorce or death of the husband. Supplementary Table 4 presents a summary of these covariates. A full list of GWAS phenotypes and their description are provided in Supplementary Table 3.

GWAS was performed using GEMMA v0.94.1 [46]. Univariate linear mixed model (LMM) as implemented in GEMMA was used to control for both population structure and genetic relatedness [46]. For each phenotype, we first removed individuals with no information on either the focal phenotype or its associated covariates. Second, we kept SNPs with maf≥ 5% for the chosen subset of individuals. Third, the standardized genetic covariance matrix was calculated from this data set and was used for LMM. Last, GWAS was run controlling for the above covariates. For continuous and count data, we provided raw phenotype data together with covariates to the program. For the binomial data, we fitted a binomial regression model using the “glm” function in R, calculated the difference between the observed odds and the odds of the fitted value, and used this residual as a GWAS phenotype. LRT *p*-values from GEMMA were used to assess significance of genetic association. *P*-values of the full and subsample sets were highly correlated for each fertility phenotype (Pearson *r* = 0.36 − 0.74 with *p* ≤ 10^−15^ for −log_10_ transformed *p*-values).

### Genomic scans for selective sweeps

To summarize genomic signatures of recent selective sweeps in Tibetans, we calculated the population branch statistic (PBS) [34] across the Tibetan genome. We used the 344 unrelated Tibetans genotyped using the Illumina OmniExpress array. For the comparison group and outgroup, we used 1KGP phase 3 CHB and CEU (CEPH Utah residents with Northern and Western European ancestry) respectively. We calculated PBS following Weir and Cockerham’s definition of pairwise *F_ST_* [90] using a custom python script for markers with maf≥ 0.05 in either Tibetans or CHB. Only female individuals from the 1KGP data were used to calculate statistics for the X chromosome. After calculating PBS, we summarized the signal for 100 kb windows sliding by 25 kb, by calculating a pseudo-binomial *p*-value *P* defined as:

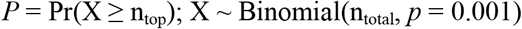

where n_top_ and n_total_ represent the number of global top 0.1% SNPs and the number of total SNPs in each window. Cutoffs for the top 0.1% PBS value were calculated separately for the autosomes and the X chromosome.

### Polygenic adaptation signals

To detect signatures of polygenic adaptation, we investigated systematic changes in allele frequencies of SNPs associated with each phenotype. For all of Tibetan GWAS phenotypes, we first took all SNPs with *p* ≤ 10^−4^ and lumped them into peaks by allowing maximum inter-SNP distance of 200 kb. Finally, we chose one SNP with the smallest association *p*-value for each peak to retrieve a set of independently associated SNPs. We also retrieved a set of SNPs associated with blood hemoglobin level (Hb) using summary statistics from a published large-scale GWAS meta-analysis [57]. For this, we first confined markers to those overlapping with our Tibetan data and applied a more stringent cutoff of *p* ≤ 10^−5^ and trimmed markers by removing one with larger association p-value from each pair of SNPs if r^2^ > 0.2 in 1KGP CEU.

After retrieving phenotype-associated SNPs with their effect size, we first compared mean frequency difference of trait-increasing alleles between Tibetans and 1KGP CHB. Following [13], we sampled 10,000 sets of random SNPs, where each set contained an equal number of SNPs as the GWAS SNPs matched one-to-one by mean minor allele frequency in bins of size 0.02. The empirical distribution of mean frequency difference of trait-increasing alleles was compared to the observed value from the GWAS SNPs and the empirical one-sided p-value was calculated as the proportion of random SNP sets with their mean allele frequency difference equal to or more extreme than the observed one.

We also looked into comprehensive signatures of polygenic adaptation using a machinery introduced by [14]. For this, we used allele frequency of 26 populations in the 1KGP phase 3 data set overlapping with the Tibetan data. We first sampled random SNPs matching each of the GWAS SNPs by minor allele frequency bin of size 0.02 in the GWAS population and by the B-value bin of size 100 (values ranging from 0 to 1,000) [91]. We sampled up to several thousands of random SNPs per GWAS SNP to obtain around 100,000 random SNPs in total. These random SNPs were used for calculating the genetic covariance matrix of populations and for generating 5,000 sets of matched random SNPs.

### Connecting selection coefficient and the statistical power to detect genotype-phenotype association

To estimate the strength of positive selection to generate difference in fertility count phenotypes large enough to be detected within a single generation, we assumed a simple additive model. That is, genotypes with 0, 1 and 2 adaptive alleles, with population frequency *p*, have the mean absolute fitness *W_0_*, *W_0_* (1+s) and *W_0_* (1+2s). Using the observed mean phenotype value, *W_m_*, we can get the per-allele effect size s *W_0_* as a function of s, *W_m_* and *p*:

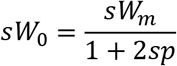

Then, the effect size was standardized to the unit of standard deviation, using the observed standard deviation of the phenotype. For the standardized effect size, which is a function of selection coefficient *s*, the statistical power to detect association was calculated using the “pwr.r.test” function in the R package “pwr”.

## Acknowledgements

Sample collection took place during the April to August 2012 with support from the National Science Foundation (1153911). Additional funding was granted through the Rockefeller Center and the Claire Garber Goodman Fund, both at Dartmouth College. Genotyping and the statistical analyses were done with support from the National Institutes of Health (1R01HL119577 to AD) and from the National Science Foundation (1153911 to CMB). We are grateful to the Nepal Health Research Council for reviewing and approving this project. We would like to thank our fieldwork assistants for their hard work and dedication to the project. In Gorkha District, they were Ang Tsering, Jangchuk Sangmo, Tinley Tsering, Tsechu Dolma, and Tsering Buti. In Mustang District, they were Kunzom Thakuri, Karma Chodron Gurung, ‘Apu’ Karma Chodron Gurung, Diki Dolkar Gurung, Yangjin Bista, Karchung Mentok Gurung, and Tashi Bista. We thank Jonathan Pritchard and Molly Przeworski for helpful comments on earlier versions of the manuscript. We are grateful to William Buikema for optimizing and performing the *EGLN1* genotyping assays and, more generally to the University of Chicago Genomics Core Facility supported by a Cancer Center Grant (P30 CA014599).

